# Kynurenic acid promotes A1 to A2 astrocyte conversion against HIV-associated neurocognitive disorder

**DOI:** 10.1101/2022.07.11.499523

**Authors:** Jingxian Lun, Yubin Li, Xuefeng Gao, Zelong Gong, Xiaoliang Chen, Jinhu Zou, Chengxing Zhou, Yuanyuan Huang, Bingliang Zhou, Pengwei Huang, Hong Cao

## Abstract

Although extensive astrocyte activation is present in the CNS of patients suffering from HIV-associated neurocognitive disorder (HAND), little is known about the contribution of reactive astrocytes to HAND neuropathology. Here, we report that the imbalance between neurotoxic (A1 astrocytes) and protective astrocytes (A2 astrocytes) occurring in HIV-1 gp120 transgenic mice impedes the resolution of inflammation in the CNS, perpetuating neuron damage and cognitive deficits. Kynurenic acid (KYNA), a tryptophan metabolite with α7 nicotinic acetylcholine receptors (α7nAChR) inhibitory property, blunts gp120-induced A1 astrocyte activation while promoting A2 astrocyte formation through blockade of α7nAChR/JAK2/STAT3 signaling activation. Meanwhile, we provide evidence that α7nAChR deletion and tryptophan administration convert astrocyte phenotypes, ultimately facilitating neuronal and cognitive improvement in the gp120tg mice. These initial and determinant findings mark a turning point in our understanding of the role of α7nAChR in gp120-mediated astrocyte activation, which opens new opportunities to control reactive astrocyte generation through KYNA and tryptophan administration as a pharmacological approach for treating HAND.

## Introduction

Despite tremendous progress in combination with antiretroviral therapy, HIV-associated neurocognitive disorder (HAND) remains a common complication in HIV-1 infected patients, especially those with improved life expectancy (Harezlak, Buchthal et al., 2011, Maschke, Kastrup et al., 2000, Tozzi, Balestra et al., 2007). One possible explanation underlying the pathogenicity of HAND is the sustained immune activation and cytotoxic effects produced by the critical HIV-1 glycoprotein 120 (gp120), leading to a chronic inflammation scenario in the CNS (He, Yang et al., 2020, Wallace, 2021). Astrocytes, the most abundant glial cells in the CNS, are functionally indispensable for maintaining neuronal survival, synaptic functions, and the blood-brain barrier (Verkhratsky & Nedergaard, 2018). The heterogeneous population of reactive astrocytes can turn into a detrimental (A1 astrocytes) or neuroprotective phenotype (A2 astrocytes) upon disease context (Escartin, Galea et al., 2021). Beyond the discrepant terminology, whether the biphasic role of reactive astrocytes in the several disorder onset and progression also holds for gp120 triggered neurodegeneration is still needed to be directly addressed (Ben Haim, Carrillo-de Sauvage et al., 2015).

Kynurenic acid (KYNA), the end-stage product of tryptophan metabolism, is primarily synthesized and liberated by cerebral astrocytes (Cervenka, Agudelo et al., 2017, Kiss, Ceresoli-Borroni et al., 2003, Moroni, Russi et al., 1988). Later studies confirmed that endogenous KYNA, at physiological concentrations, acts predominantly as a non-competitive antagonist of α7 nicotinic acetylcholine receptors (α7nAChR) (Hilmas, Pereira et al., 2001). α7nAChR are abundant in the hippocampus, notably in the astrocytes, participating in neuronal synaptic transmission, memory, and cognition processes (Bertrand, Lee et al., 2015, Shen & Yakel, 2012). Additionally, α7nAChR stimulation recruits Janus kinases 2 (JAK2)/signal transducer and activator transcription 3 (STAT3), a canonical molecule controlling astrocyte reactive transformation (de Jonge, van der Zanden et al., 2005, Giovannoni & Quintana, 2020). Given that α7nAChR activation leads to the recruitment of JAK2/STAT3 to its receptor complex, α7nAChR inhibition through KYNA may offer a plausible approach to ameliorate reactive astrocyte generation under gp120 insult.

Here, we took advantage of primary astrocyte cultures, gp120 transgenic mice (gp120tg mice), α7nAChR knockout mice (α7^-/-^ mice), and α7^-/-^gp120tg mice and aimed to investigate the effect of KYNA on functional transformation in gp120-induced reactive astrocytes. We tested the hypothesis that inhibition of α7nAChR mediated JAK2/STAT3 signaling pathway may block neurotoxic astrocyte activation driven by rising levels of gp120. Finally, we examined whether tryptophan, a source of endogenous KYNA, could improve histopathologic changes in the brain of gp120tg mice via altering the reactive states of astrocytes.

## Results

### Progressive neurotoxic astrocyte activation in gp120tg mice with age-related neurological deficits

The gp120tg mice featuring the expression of viral gp120 protein in CNS astrocytes under the control of a modified *GFAP* promoter is a surrogate model for HAND (Toggas, Masliah et al., 1994). Initially, we analyzed histological changes by H&E and Nissl staining for the hippocampal subregions, where gp120tg mice displayed essential pyknosis in cells and visible neuronal loss (Figure EV1A). Meanwhile, the gp120tg mice exhibited an age-dependent loss of neuronal nuclei (NeuN), microtube-associated protein 2 (MAP-2), and synaptophysin in the hippocampus and cortex (Fig. 1A, Figure EV1B). WT mice consistently outperformed 12-month-old gp120tg mice on each training day during the Morris water maze (MWM) test (Fig. 1B-C). In the probe trial, gp120tg mice showed a concentric swimming pattern within proximity of the wall while spending less time in the targeted quadrant (Fig. 1D-E). Furthermore, cognitive performance confirmed age-dependent memory impairments in 3-, 6-, 12-, and 24-month-old gp120tg mice, suggesting that gp120 expression causes progressive neuronal destruction accompanied by robust cognitive deficits in mice (Figure EV1C-F). Regarding emotionality, 12-month-old gp120tg mice exhibited decreased dwell in the center of the arena, displaying an anxiety-like behavior in the open field test (Figure EV1G-J). Complement component 3 (C3) markedly enriched in neurotoxic reactive astrocytes from patients with neurodegenerative disorders (Liddelow, Guttenplan et al., 2017). Indeed, the C3^+^/GFAP^+^ cells in the hippocampal subregion of 12-month-old gp120tg mice were already evident and dramatically increased during the disease progression (Fig. 1F). Strikingly, we observed a significant age-dependent increase in A1-specific genes and proinflammatory cytokines in the hippocampus, where the A1 astrocyte activation preceded frank neuronal injury at the early age of gp120tg mice (Fig. 1G and I). These transcriptomic features were associated with moderate upregulations of A2-specific genes and neurotrophic factors in gp120tg mice (Fig. 1H and I). Together, our results suggest that gp120-elicited pathological progression in CNS was partially neurotoxic reactive astrocyte inducing.

**Figure 1.**
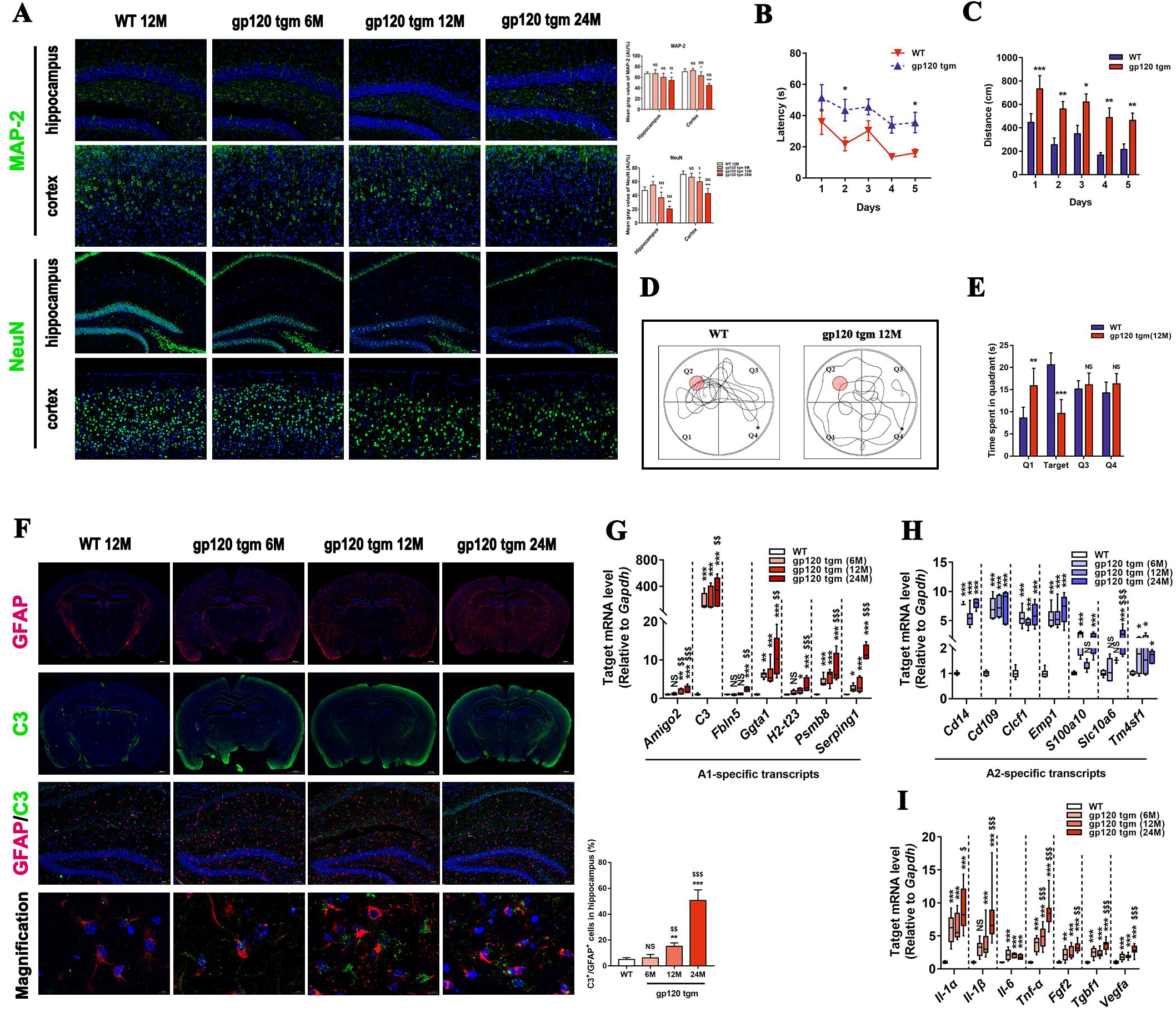
HIV-1 gp120 expression induces progressive neuropathology and A1 astrocyte activation in mice. A Immunolabeling of neuronal cells (NeuN) and dendritic structures (MAP-2) in the hippocampus and cortex (*n*=5 mice per group, scale bar: 100 μm). B-E The 12-month-old gp120tg mice and age-matched WT mice were subjected to the MWM test to assess their long-term spatial memory. Latency (B) and distance (C) to locate the submerged platform during 5 training days. (D) Searching strategies for the removed platform in the probe trial. Black dots depicted the initial position, and white dots represented the final position. (E) Time spent in the target quadrant in the retrieval test (*n* = 10 mice per group). F Immunofluorescence staining against GFAP (red) and C3 (green) in the whole brain sections (scale bar: 1000 μm) and the hippocampus (scale bar: 100 μm). High-magnification images showed co-localization of GFAP and C3 (scale bar: 10 μm). *n* = 5 mice per group. G-I The mRNA expression profiles of A1-specific genes (G), A2 phenotype markers (H), and cytokines (I) in the hippocampus of mice (*n* = 5 mice per group). Data information: In (A-E), data are presented as mean ± SD, Student’s *t* test, two-tailed. In (F), data are analyzed by One-way ANOVA. In (G-I), values are shown in the box and whisker plot where the line in the box corresponds to the median. One-way ANOVA. \**P* < 0.05, \*\**P* < 0.01, \*\*\**P* < 0.001, NS *P* > 0.05 (not significant) compared to WT mice. $*P* < 0.05, $$*P*<0.01, ^$$$^*P* < 0.001 compared to 6-month-old gp120tg mice.

### KYNA controls the reactive transformation in gp120-induced astrocytes

KYNA, long known as a neuroprotective modulator, is mainly synthesized in astrocytes within the CNS (Kiss et al., 2003). As expected, the immunoreactivity of C3 and GFAP induced by gp120 waned upon KYNA administration (Fig. 2A-B, Figure EV2A). The gp120-induced astrocytes displayed a reversed expression profiles enriched for mRNA encoding the A2 phenotype, not the A1 phenotype, following KYNA incubation (Fig. 2C-D). Furthermore, KYNA co-treatment substantially suppressed pro-inflammatory cytokines released in the supernatant compared to the gp120 group, with up to approximately 50% inhibition at 25 μM KYNA concentration (Fig. 2E). On the contrary, KYNA stimulated the production of other cytokines, including brain-derived neurotrophic factor (BDNF), glial cell line-derived neurotrophic factor (GDNF), and nerve growth factor (NGF), which were at low levels after gp120 infection (Fig. 2F). Since KYNA acts as a negative allosteric regulator of α7nAChR, we sought to characterize what downstream events that followed blockage of α7nAChR altered in gp120-induced astrocytes under KYNA treatment. The enhancement in fluorescence intensity on cell membranes indicated that α7nAChR was abundant in astrocytes following the challenge with high-dose gp120 (Figure EV2B-C). The inhibitory effect of KYNA on expression of α7nAChR, JAK2, STAT3, and phosphorylated counterparts in gp120-treated astrocytes was efficient (Fig. 2G, Figure EV2D-F). Collectively, we suggest that KYNA blunts the activation of neurotoxic astrocytes provoked by gp120 infection while promoting the formation of neuroprotective astrocytes, presumably due to the α7nAChR/JAK2/STAT3 signaling inhibition.

**Figure 2.**
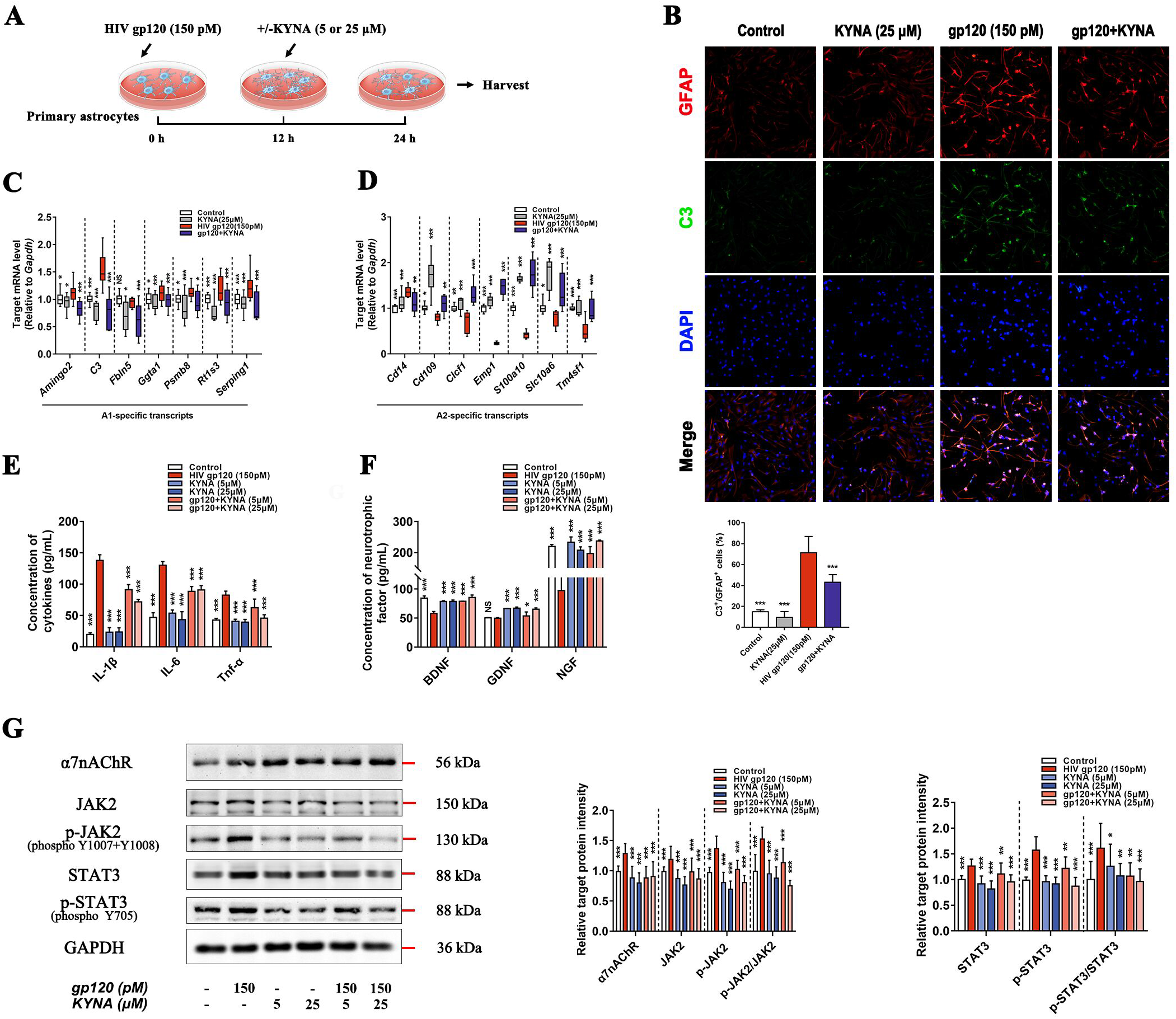
KYNA promotes the switching of A1 to A2 astrocytes. A Paradigm depicting KYNA treatment. B Double immunofluorescence images showing the expression of GFAP (red), C3 (green), and nucleus (blue) in primary astrocytes (scale bar: 50 μm). C, D Relative gene expression of A1 phenotype (C) and A2 phenotype (D) in primary astrocytes. E, F Quantification of (E) Il-1β, Il-6, Tnf-α, (F) BDNF, GDNF, and NGF levels in cultured supernatant. G Protein expression of the α7nAChR/JAK2/p-JAK2/STAT3/p-STAT3 relative to the housekeeping protein GAPDH. Data information: In (B, E-G), data are presented as mean ± SD. In (C-D), values are shown in the box and whisker plot where the line in the box corresponds to the median. Throughout, *P* values are calculated using one-way ANOVA with *n* = 5 cultures per group. \**P* < 0.05, \*\**P* < 0.01, \*\*\**P* < 0.001, NS *P* > 0.05 (not significant) compared to 150 pM HIV-1 gp120 treated group.

### Blockade of α7nAChR promotes the conversion of A1 astrocytes to A2 astrocytes after gp120 infection

Methyllycacontitine (MLA), a selective antagonist at α7nAChR, with escalating doses of KYNA, pronouncedly reduced the expression of α7nAChR/JAK2/p-JAK2/STAT3/p-STAT3 in gp120-induced astrocytes as comparisons to mere KYNA treated groups (Fig. 3A-B). Meanwhile, MLA potentiated the inhibitory effects of KYNA on the expression of C3 and GFAP elevated by gp120 in astrocytes (Fig. 3C-E). The decreased expression of A1-specific genes concomitant with the upregulation of A2-specific gene expression was observed in the MLA-treated astrocytes (Fig. 3F-G). KYNA promoted the heterogeneous effect of MLA on phenotype transformation of reactive astrocytes, indicating that α7nAChR is essential for astrocyte conversion (Fig. 3F-G). We next wondered whether the phenotypic transformation mediated by α7nAChR inhibition might, in turn, reverse the functions of astrocytes. Indeed, remarkable decreases in inflammatory cytokines paralleled a slight increase of neurotrophic factors during sustained blockage of α7nAChR in gp120-induced astrocytes (Fig. 3H-I). Even though a fraction of A1-specific genes was increased in the astrocytes following nicotine (a nAChRs activator) or PNU-282987 (an α7nAChR agonist) pretreatment, KYNA still slightly attenuated A1 astrocyte responses while dramatically boosting A2 astrocyte responses (Figure EV3). Overall, we conclude that KYNA exerts its reversed impact on the phenotypic formation and function of gp120-induced astrocytes, to a large extent, via inhibition of the α7nAChR/JAK2/STAT3 signaling pathway.

**Figure 3.**
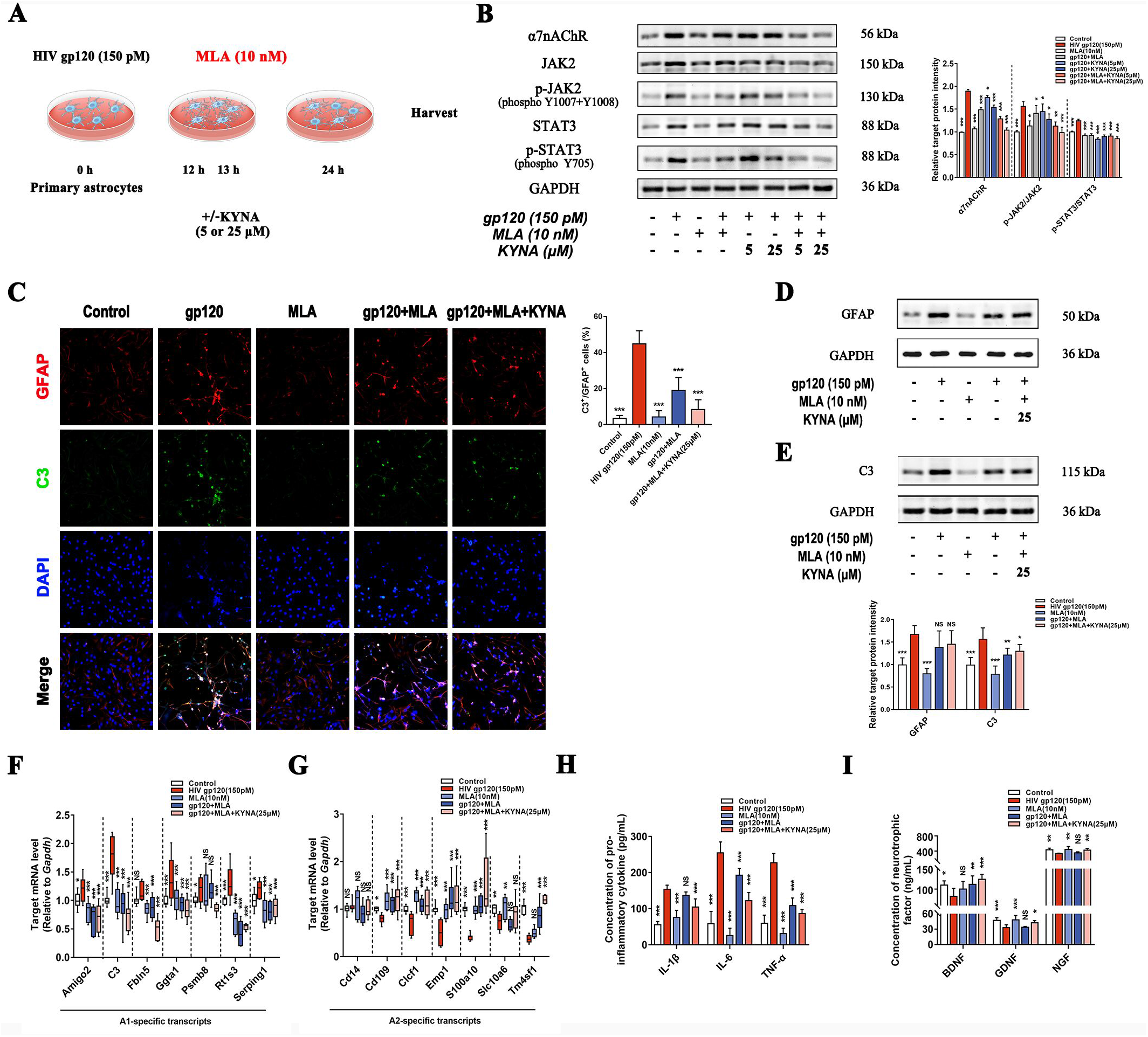
KYNA synergies with MLA against gp120-induced neurotoxic transformation in astrocytes. A The detailed paradigm for MLA treatment. B Western blot demonstrating the α7nAChR, JAK2, p-JAK2, STAT3, and p-STAT3 protein levels in the astrocytes. C Images of GFAP (red) and C3 (green) positive cells (scale bar: 50 μm). D, E Immunoblot analysis of the cell lysates presenting the expression of GFAP (D) and C3 (E) proteins. F, G Quantification of the expression of A1 specific genes (F) and A2 specific genes (G). H, I Protein levels of Il-1β, Il-6, Tnf-α (H), BDNF, GDNF, and NGF (I). Data information: In (B-E, H-I), data are shown as the mean ± SD. In (F-G), values are shown in the box and whisker plot where the line in the box corresponds to the median. one-way ANOVA, *n* = 5 cultures per group. \**P* < 0.05, \*\**P* < 0.01, \*\*\**P* < 0.001, NS *P* > 0.05 (not significant) compared to 150 pM HIV-1 gp120 treated group.

### Involvement of JAK2/STAT3 signaling in the phenotypic switch in reactive astrocytes

We then pretreated astrocytes with AG490, a well-characterized inhibitor for JAK2, to mimic JAK2/STAT3 inhibition via KYNA (Fig. 4A). In AG490-stimulated astrocytes with reduced STAT3 and p-STAT3 expression, the decrease in p-STAT3/STAT3 ratio remained pronounced upon KYNA interference (Fig. 4B-C). Furthermore, KYNA yielded a remarkable decrease in the expression of A1-specific genes and a slight upregulation of A2-specific genes in astrocytes, even co-applied with the AG490 (Fig. 4D-E). Strikingly, AG490 and KYNA showed a reciprocal relationship in attenuating the release of proinflammatory cytokines imposed by gp120 in astrocytes (Fig. 4F-G). Later experiments showed that inhibition of JAK2/STAT3 responses greatly reversed the expressions of neurotrophic factors limited in the gp120-induced astrocytes (Fig. 4H). Collectively, our data suggest that JAK2/STAT3 signaling inhibition is critical for KYNA to blunt the neurotoxic responses while promoting neuroprotective responses in gp120 infected astrocytes.

**Figure 4.**
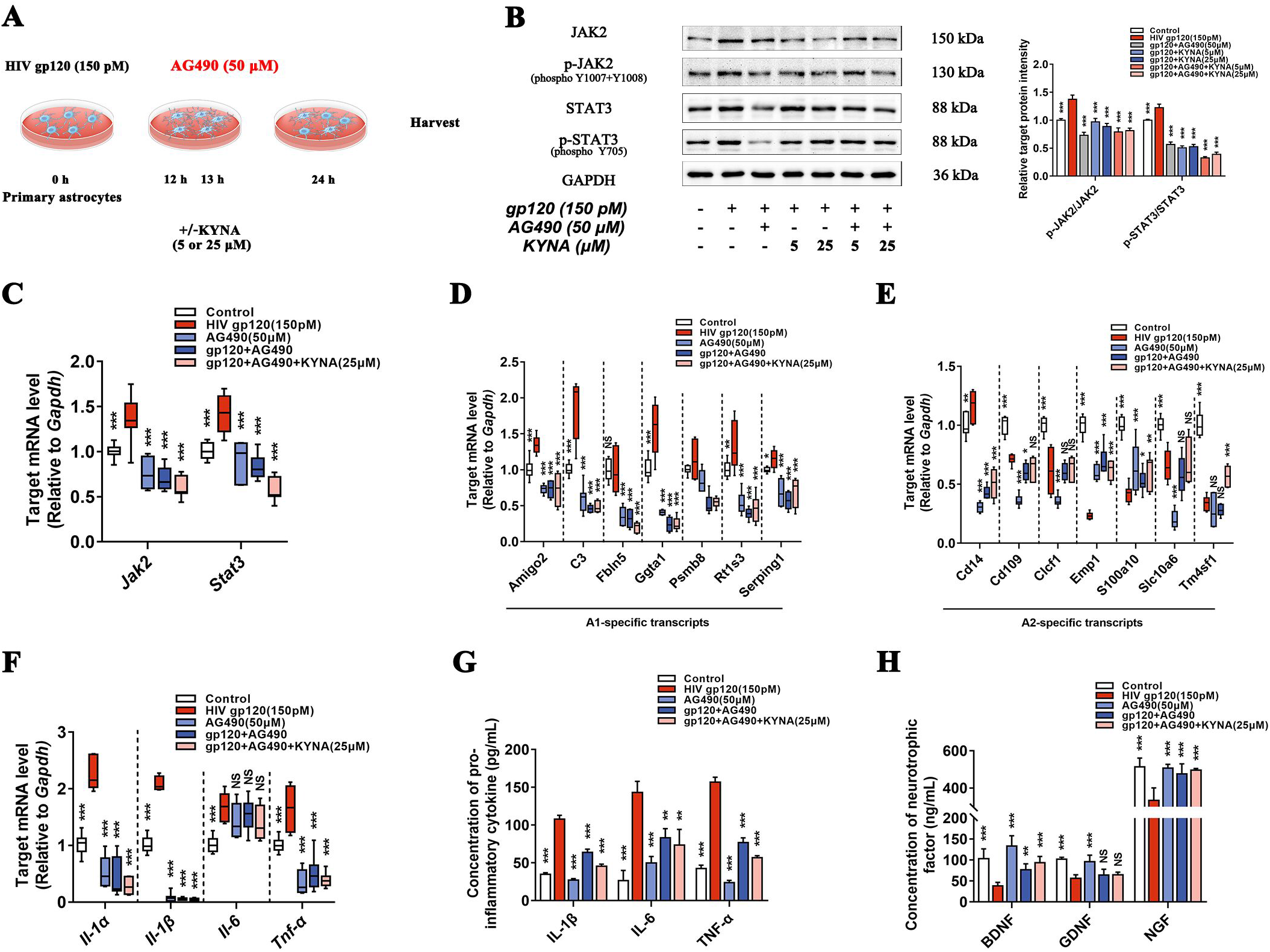
Inhibition of neurotoxic responses in gp120-stimulated astrocytes by targeting the JAK2/STAT3 signaling pathway. A Experimental paradigm for AG490 treatment. B Immunoblots for JAK2/p-JAK2/STAT3/p-STAT3 protein levels in stimulated astrocytes. C The mRNA levels of *Jak2* and *Stat3*. D-F Quantification of A1-specific gene (D), A2-specific gene expressions (E), and cytokines (F) in each treated group. G, H The protein levels of Il-1β, Il-6, Tnf-α (G), BDNF, GDNF, and NGF (H) in astrocyte culture supernatants as measured by ELISA. Data information: In (B, G-H), data present mean ± SD. In (C-F), values are shown in the box and whisker plot where the line in the box corresponds to the median. *P* values were calculated using one-way ANOVA with *n* = 5 biologically independent cell cultures. \**P* < 0.05, \*\**P* < 0.01, \*\*\**P* < 0.001, NS *P* > 0.05 (not significant) compared to 150 pM HIV-1 gp120 treated group.

### α7nAChR knockout rescues cognitive impairment in gp120tg mice via modulation of reactive astrocyte transformation

To pinpoint the role of α7nAChR, we employed a knockout mice model with a null mutation of the *Chrna7* gene, referred to as α7nAChR knockout mice (α7^-/-^ mice). The changes of A1 and A2 specific genes in the hippocampus of α7^-/-^ mice were negligible compared to age-match WT mice (Fig. 5A-B), indicating that the *Chrna7* gene deletion alone did not evoke the production of reactive astrocytes. To address whether α7nAChR expression is a prerequisite for gp120-elicited phenotypic changes in astrocytes, we mated the α7^-/-^ mice with gp120tg mice line to generate a cross-breed mice model, α7^-/-^gp120tg mice which expressed gp120 with a concomitant α7nAChR deficiency. As expected, the α7^-/-^gp120tg mice exhibited virtually down-regulated A1-specific genes while sustaining A2-specific gene responses, which resulted in a weaker immunoreactivity of GFAP and C3 in the hippocampus (Fig. 5C-E). In line with the limited expression of inflammatory cytokines in α7^-/-^ mice, *Chrna7* deletion robustly reduced the production of proinflammatory cytokines whereas moderately increasing *Tgfb-1* mRNA levels in the gp120tg mice (Fig. 5F-G). Simultaneously, the α7nAChR knockout leads to the truncation of JAK2/STAT3 signal responses in the hippocampus of α7^-/-^gp120tg mice (Fig. 5H). Furthermore, with data collapsed across the 5 training days in the MWM test, α7^-/-^gp120tg mice slightly outperformed gp120tg mice each day during the training session (Fig. 5I-J). The gp120tg mice exhibited wall-hugging behavior in the probe trials, whereas the α7^-/-^gp120tg mice used a focal search strategy for the removed platform showing a clear preference for the target quadrant (Fig. 5K-M). We also observed decreased anxiety-like behavior of the α7^-/-^gp120tg mice in the open field apparatus (Fig. 7N). Together, our data reinforce that ablation of α7nAChR is necessary and sufficient to shield mice from gp120-induced neurotoxicity and cognitive declines.

**Figure 5.**
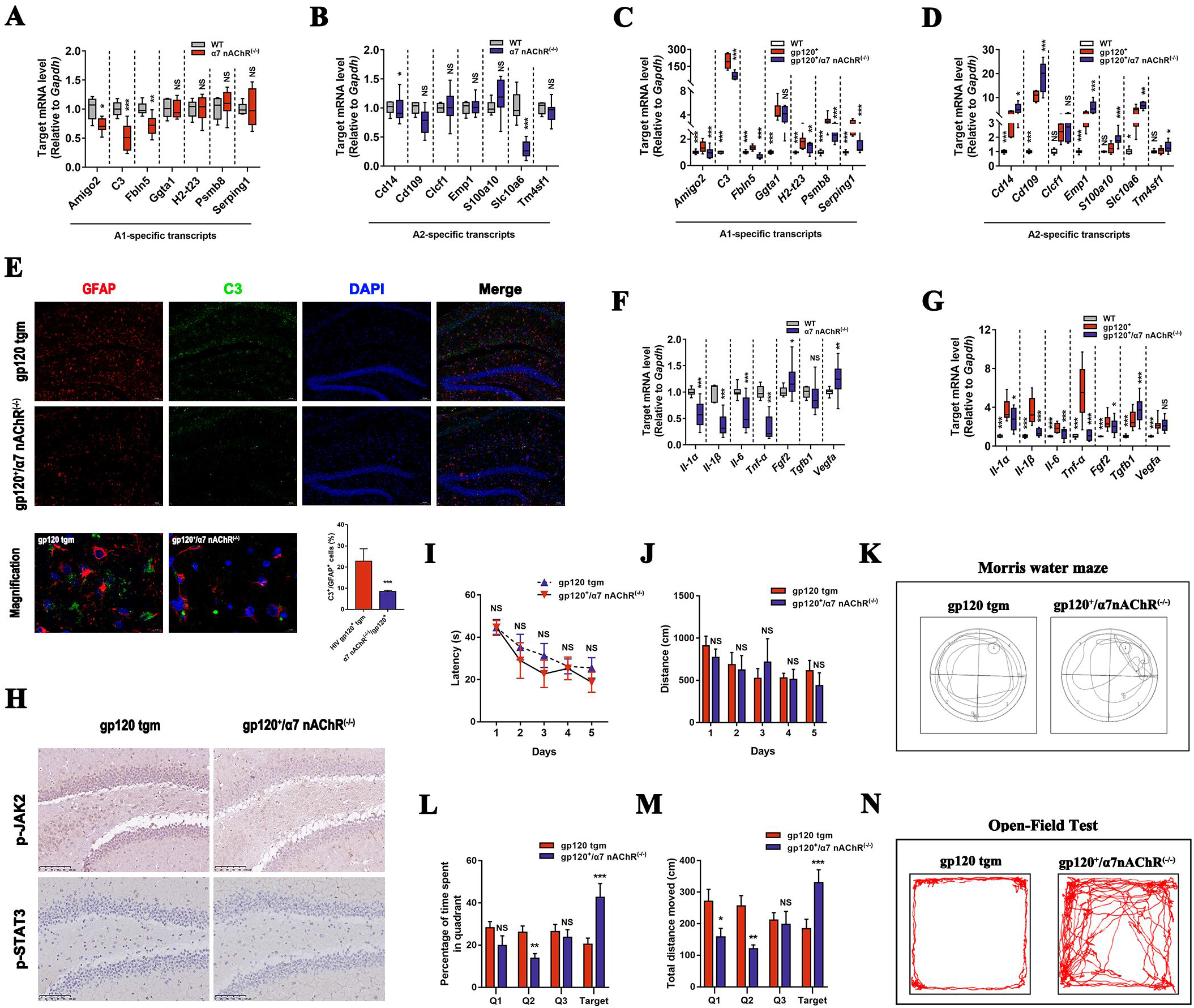
Ablation of the Chrna7 gene in gp120tg mice restores the neuroprotective capacity of reactive astrocytes. A, B The expression of A1-specific genes (A) and A2-specific genes (B) in the hippocampus from 12-month-old α7^-/-^ mice and age-matched WT mice (*n* = 6 mice per group). C-D The mRNA levels of A1-specific genes (C) and A2-specific genes (D) in 12-month-old gp120tg mice and age-matched α7^-/-^gp120tg mice (*n* = 5 mice per group). E The immunoreactivity of GFAP and C3 proteins in the hippocampus (scale bar: 100 μm). High-magnification images showed co-labeling of GFAP and C3 (scale bar: 10 μm). *n* = 5 mice per group. F The mRNA levels of *Il-1α, Il-1β, Il-6, Tnf-α, Fgf-2, Tgfb-1*, and *Vegf-a* in the hippocampus of α7^-/-^ mice and WT mice (*n* = 6 mice per group). G The mRNA levels of cytokines among WT mice, gp120tg mice, and α7^-/-^gp120tg mice (*n* = 5 mice per group). H Representative images of immunohistochemistry staining for p-JAK2 and p-STAT3 proteins on brain sections from the hippocampus subregion of the gp120tg mice and α7^-/-^gp120tg mice (scale bar: 100 μm). *n* = 5 mice per group. I-M The gp120tg mice and α7^-/-^gp120tg mice were subjected to the Morris water maze test. The discrepancy in latency (I) and distance (J) between gp120tg and α7^-/-^gp120tg mice during acquisition trails. (K) Search strategies for the removed platform in the retrieval test. Percentage of time spent in the different zones (L) and total distance (M) in the probe trials (*n* = 10 mice per group). N Characteristic navigation patterns of mice in the open field test (*n* = 10 mice per group). Data information: In (A-D, F-G), values are shown in the box and whisker plot where the line in the box corresponds to the median. In (I-M), data present mean ± SD. In (A-B, F, I-M), *P* values are analyzed by unpaired Student’s t test (two-tailed). In (C-D, G), *P* values are analyzed by one-way ANOVA. \**P* < 0.05, \*\**P* < 0.01, \*\*\**P* < 0.001, NS *P* > 0.05 (not significant).

**Figure 6.**
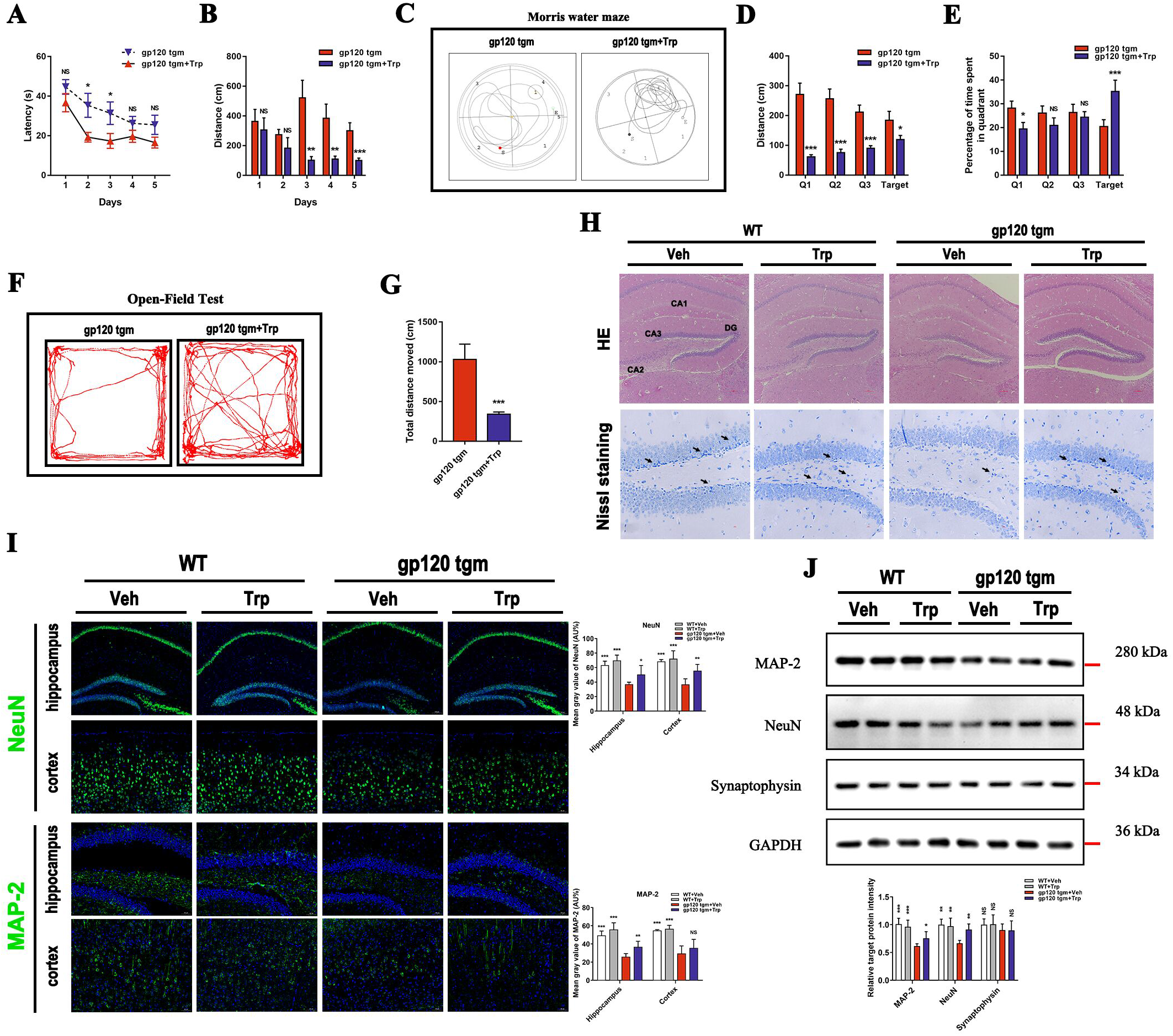
Tryptophan affords protection against cognitive declines and aberrant neurons in gp120tg mice. A-E The gp120tg mice were allowed to drink water containing 0.1% tryptophan for 1 month, then were subject to the Morris Water Maze test. Latency to reach the hidden platform (A) and cumulative swim distance (B) between the two groups during the acquisition phase. (C) Plots showing the different searching strategies of gp120tg mice and tryptophan-fed gp120tg mice in probe trials. Distance (D) and percentage of time spent (E) in the targeted quadrant in the probe trial (*n* = 10 mice per group). F-G Behavior (F) and cumulative distance (G) of gp120tg mice following tryptophan administration during the open field test. H Brain sections stained with hematoxylin and eosin (scale bar: 100 μm). Magnification of Nissl staining section in the hippocampus subregion. Black arrows indicated the Nissl bodies (scale bar: 10 μm, *n* = 5 mice per group). I Immunofluorescence showed the distribution of the NeuN and MAP-2 in the hippocampus and cortex among different groups (scale bar: 100 μm, *n* = 5 mice per group). J The protein levels of NeuN, MAP-2, and Synaptophysin in the cortex and hippocampus (*n* = 5 mice per group). Data information: Throughout, data present mean ± SD. In (A-G), *P* values are analyzed by unpaired Student’s *t* test (two-tailed). In (I-J), data are analyzed by one-way ANOVA. \**P* < 0.05, \*\**P* < 0.01, \*\*\**P* < 0.001, NS *P* > 0.05 (not significant) compared to 12-month-old gp120tg mice treated with PBS.

**Figure 7.**
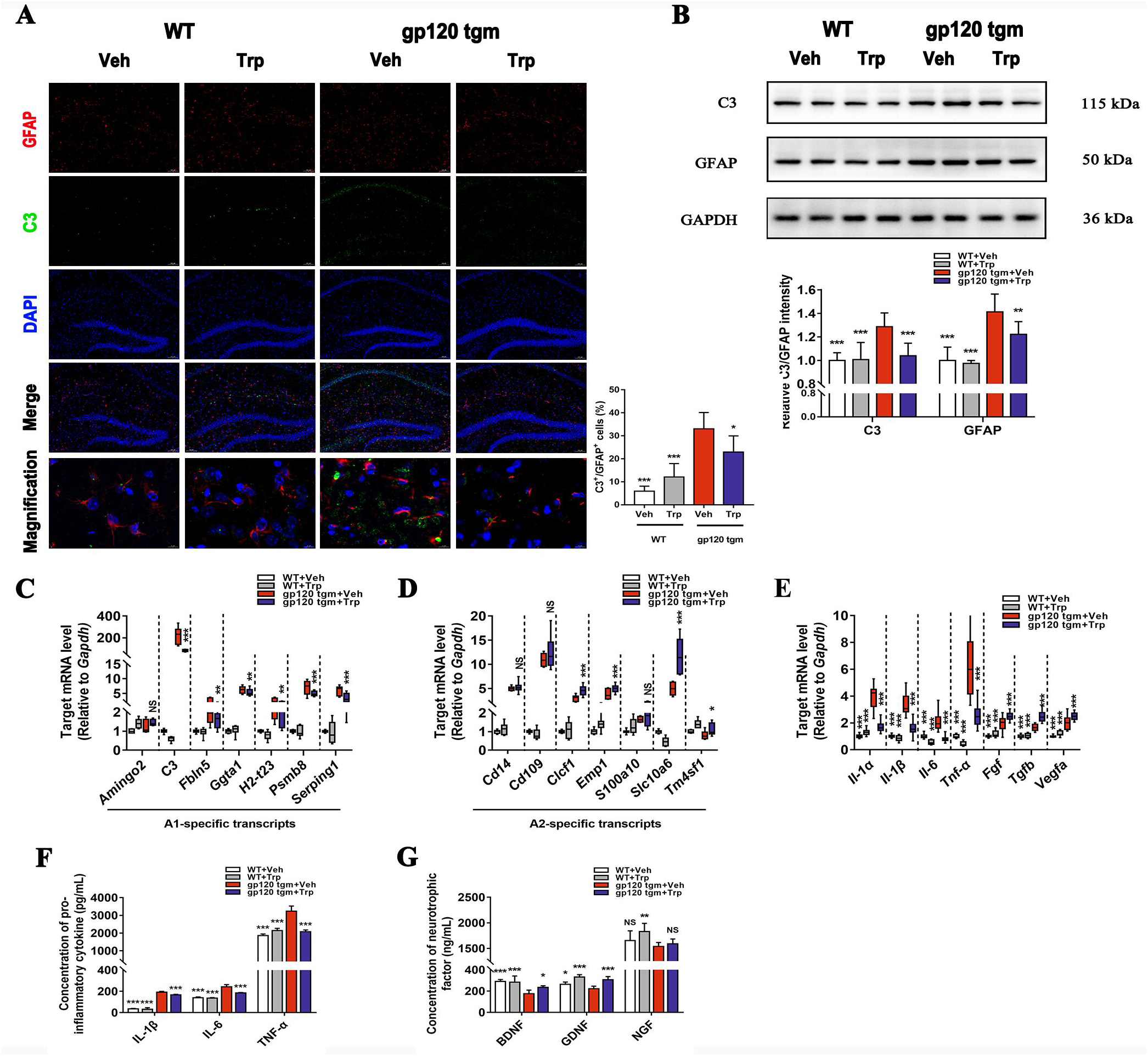
Tryptophan alleviates neurotoxic responses in gp120tg mice while preserving the neuroprotective effects of astrocytes. A Colocalization of GFAP (red) and C3 (green) cells in the hippocampus from mice treated with PBS or tryptophan (scale bar: 100 μm). High-magnification images showed co-staining with GFAP and C3 (scale bar: 10 μm). B GFAP and C3 protein levels in the brain lysates as determined by Western blot. C-E Alterations of A1-specific genes (C), A2-specific genes (D), and cytokines (E) in the hippocampus of mice following tryptophan administration for 1 month. F, G Serum levels of Il-1β, Il-6, Tnf-α (F), BDNF, GDNF, and NGF (G) as measured by ELISA. Data information: In (A-B, F-G), data present mean ± SD. In (C-E), values are shown in the box and whisker plot where the line in the box corresponds to the median. Throughout, representative data of three independent experiments with *n* = 5 mice per group. \**P* < 0.05, \*\**P* < 0.01, \*\*\**P* < 0.001, NS *P* > 0.05 (not significant) compared to 12-month-old gp120tg mice treated with PBS.

### Tryptophan improves cognitive deficits in gp120tg mice through inhibition of neuronal injury

We designed to reintroduce dietary tryptophan into mice in case to elevate the endogenous KYNA levels in the CNS to investigate what extent tryptophan can alter widespread neuropathology in 12-month-old gp120tg mice. Mice receiving the tryptophan diet spent remarkably shorter time and distance finding the hidden platform than gp120tg mice did on the 1^st^ day onwards, as shown in the acquisition phase of the MWM test (Fig. 6A-B).

Furthermore, tryptophan-fed mice showed a focal track targeting the removed platformed in the probe trials (Fig. 6C), which resulted in the prolonged dwell in the targeted quadrant (Fig. 6D-E). Notably, gp120tg mice were active with high avoidance of the center, whereas mice supplemented with tryptophan showed a relatively low avoidance of the central area in the open field test (Fig. 6F-G). In addition, vacuolization and neuronal loss in the hippocampus appeared at a much lower degree in tryptophan-received mice (Fig. 6H). The increased expression of NeuN, MAP-2 and synaptophysin protein levels in the tryptophan-fed gp120tg mice was significant (Fig. 6I-J), suggesting that tryptophan treatment rescues already established neuronal abnormalities in gp120tg mice.

### Tryptophan attenuates neurotoxic responses in gp120tg mice via promoting phenotypic reversion in astrocytes

Consistent with our previous findings, we assumed that the neuropathological improvement in tryptophan-fed mice is also associated with the altered reactive states of astrocytes. Indeed, tryptophan robustly blocked the overactivation of C3^+^/GFAP^+^ cells in the hippocampus of gp120tg mice (Fig. 7A-B). This result was mirrored by remarkably decreased C3 and GFAP protein levels in tryptophan-fed gp120tg mice (Fig. 7B). The transcriptomic responses of A1 genes were milder in the hippocampus of gp120tg mice following tryptophan administration (Fig. 7C). Conversely, the vast majority of A2-specific genes were remarkable induction in the tryptophan-received mice (Fig. 7D), suggesting that tryptophan reverses the reactive status of astrocytes as potently as KYNA. Finally, to assess whether the tryptophan introduction shapes the inflammatory microenvironment in gp120tg mice, we monitored the cytokine levels of gp120tg mice on the tryptophan diet. As expected, tryptophan counteracted the trend toward higher proinflammatory cytokine expression in protein and mRNA levels in gp120tg mice (Fig. 7E-F). Meanwhile, mice receiving tryptophan exhibited a subtle yet significant increase in the neurotrophic factors compared to gp120tg mice (Fig. 7E and G). Altogether, our data suggest that tryptophan is protective against neurotoxic responses in gp120tg mice, which is large-extent related to the phenotypic transformation of reactive astrocytes.

## Discussion

Mechanisms of ongoing neurodegeneration in HIV-infected patients remain elusive, as HIV-1 cannot directly damage neurons, whereas widespread neuronal damage is a pathological hallmark in HAND (Everall, Luthert et al., 1993). Intriguingly, resting astrocytes morphed into A1 astrocytes in response to LPS stimuli, whereas astrocytes undergo distinct gene profiles in the ischemia mice model, and this type of A2 astrocytes is putatively to be neuroprotective (Zamanian, Xu et al., 2012). Since this seminal finding, researches on astrocyte heterogeneity indicate that A1 astrocytes contribute to the progression of several neurodegenerative disorders (Joshi, Minhas et al., 2019, Liddelow et al., 2017, Yun, Kam et al., 2018).

Here, we found that HIV-1 shed protein gp120 evoked moderate astrocyte activation in mice even at a young age when overt neuropathology in the hippocampus is absent, indicating the occurrence of reactive astrocytes is causative, preceding neuronal injury. Subsequently, neurotoxic reactive astrocytes and neuronal losses were on dramatic rises causing obvious memory impairment in aged gp120tg mice. This data is reminiscent of previous studies that aging increases the sensitivity of mice to cognitive impairment and executive dysfunction following central gp120 insults postulated due to the prolonged cytokine response within the CNS (Abraham, Jang et al., 2008, Sparkman, Buchanan et al., 2019). Meanwhile, aging induces up-regulation of a cassette of A1 and A2 specific genes in mice (Clarke, Liddelow et al., 2018), consistent with the increased gene expressions of reactive astrocytes at dual states in aged gp120tg mice, indicating that A2-astrocyte activation may be a compensatory mechanism for limitation of the extensive toxic effects. Previous studies have shown that Il-1α, Tnf, and C1q, driven by the activated microglia, promote neurotoxic astrocyte formation (Liddelow et al., 2017). Consequently, Il-1α, Il-1β, and Tnf-α act back to the gp120-stimulated reactive astrocytes, which engenders a positive feedback loop that amplifies the neurotoxic response (Hennessy, Griffin É et al., 2015), explaining why the A1-response is much fierce than A2-response in both transgenic mice and primary astrocytes.

A long-standing question concerning astrocyte activation triggered by the sterile inflammatory insults is whether neurotoxic reactive astrocytes can retrospectively transform to a physiological or neuroprotective state. We demonstrated that KYNA reverts A1 to A2 phenotype through inhibiting gp120-primed α7nAChR/JAK2/STAT3 signaling cascades in primary rat astrocytes, laying a consolidation for the therapeutic efficacy of KYNA. Endogenous KYNA has neuroinhibitory properties attributed to its action as an antagonist of NMDA receptors and α7nAChR (Alkondon, Pereira et al., 2004). Functional α7nAChR up-regulation is not only found in the gp120tg mice but also the HIV-infected subjects (Ballester, Capó-Vélez et al., 2012, Delgado-Vélez, Báez-Pagán et al., 2015), pointing towards a potential role of wide-expressed α7nAChR in gp120-induced astrocytes. Moreover, Mecamylamine, a non-competitive antagonist of nAChR, strongly abolished nicotine-induced morphological changes in astrocytes, supporting the role of α7nAChR in this process (Aryal, Fu et al., 2021). After pharmacological truncation or gene deletion, both KYNA and MLA, let alone the α7nAChR deletion, dramatically reversed the reactive state and function of gp120-induced astrocytes, demonstrating that α7nAChR inhibition confers astrocytes with phenotypic transformation capacities.

To clarify the downstream events in the phenotypic conversion of astrocytes upon KYNA-mediated α7nAChR blockage, we utilize AG490 to pinpoint the JAK2/STAT3 signaling pathway, as STAT3 is highly involved in the formation of reactive astrocytes (Escartin et al., 2021). We found that AG490 and AG490-KYNA co-incubation dramatically downregulates gp120-induced A1-astrocyte formation via JAK2/STAT3 signaling inhibition, although the synergy effect seems negligible. Indeed, under several neuropathological conditions seen in previous studies, negative regulation of STAT3 signaling cascades prevents GFAP, a pivotal protein controlling the shape and movement of astrocytes, and C3 protein activation (Huang, Wang et al., 2016, Okada, Nakamura et al., 2006, Reichenbach, Delekate et al., 2019). The outcome of STAT3 positive astrocytes to disease is intricate because they are beneficial in the healing process, limiting myelin injury and structural synaptic plasticity (Nobuta, Ghiani et al., 2012, Okada et al., 2006, Tyzack, Sitnikov et al., 2014). However, they are also associated with neuropathic pain maintenance, blood-brain barrier dysfunction, iron-activated neuronal apoptosis, brain metastasis colonization, and accumulating β-amyloid levels (Huang et al., 2016, Priego, Zhu et al., 2018, Reichenbach et al., 2019, Tsuda, Kohro et al., 2011, You, Yan et al., 2017). Given the multifactorial nature of astrocytes, it should be prudent to disclose the role of the JAK2/STAT3 signaling pathway in the controlling fine-tuned process of astrocyte phenotype switch. Taken as a whole, due to disruptions of α7nAChR-mediated JAK2/STAT3 signaling transduction, KYNA administration resumes neuroprotective potentials of gp120-induced reactive astrocytes.

Previous studies have revealed a shift towards increased KYNA synthesis following inhibition of the primary route for tryptophan catabolism. Meanwhile, this metabolic alteration associated with the increased KYNA levels is neuroprotective in HD flies and Alzheimer’s disease mouse model (Breda, Sathyasaikumar et al., 2016, Campesan, Green et al., 2011, Zwilling, Huang et al., 2011). For mimicking the elevated levels of KYNA in CNS of gp120tg mice, we utilize the endogenous synthesis strategies to convert dietary tryptophan to KYNA, as KYNA cannot traffic across the blood-brain barrier due to its polar structure (Fukui, Schwarcz et al., 1991). Consistent with the reversed capacity of KYNA, we further demonstrated that tryptophan administration dramatically impedes A1-astrocyte formation while promoting A2-astrocyte formation in gp120tg mice, underlying the protective mechanism of tryptophan and its metabolites in neuronal damage and cognitive decline.

Collectively, we interrogated the dual role of reactive astrocytes to unravel the protective mechanism of KYNA and tryptophan under gp120 insults. Our findings highlight that α7nAChR inhibition facilitates the A1-to-A2 astrocyte transformation, which may contribute to the therapeutic interventions for HAND.

## Materials and Methods

### Primary cell culture and drug treatments

Primary cultures of astrocytes from the rat cortex were established and maintained as previously published protocol (Schildge, Bohrer et al., 2013). Briefly, cortical astrocytes were dissected from SD neonatal rats (1-3 days) with the aseptic operation and digested by 0.25% trypsin (Gibco, USA) at 37°C for 20 min. The cell suspension was dispersed through the cell strainer (75 μm pore size, Bioland, China) and seeded into the poly-D-lysine coated T-75 flasks (Corning, USA) with DMEM/F12 (Gibco, USA) supplemented with 10% FBS, 1% penicillin/streptomycin. After 10 days of incubation, the cells were purified in a shaking incubator (250 r/min) at 37°C for 24 h, and astrocytes were cultured at desired densities in 6-well plates. The cells were identified as >98% pure astrocytes by Cy3-conjugated GFAP immunostaining (at 1:400 dilution, Abcam, UK). Cultured astrocytes were treated with 150 pM HIV-1 gp120 CM (ProSpec-Tany TechnoGene Ltd., Israel) for 12 h before application of 5 or 25 μM KYNA (Sigma-Aldrich, USA) for a further 12 h. In addition, gp120-induced (150 pM, 11 h) astrocytes were pretreated for 1 h with media in the presence or absence of 10 nM methyllycacontitine (MLA, MCE, USA) or 50 μM AG490 (Selleckchem, USA) followed by a 12 h co-incubation period with KYNA.

### Total RNA isolation and quantitative reverse transcription PCR (RT-qPCR) analysis

According to the manufacturer’s instructions, total RNA was extracted from samples using RNAiso Plus Reagent. Reverse transcription was performed following the protocol of the PrimeScript™ RT Master Mix kit. Then, cDNA was run for quantitative PCR using TB Green^®^ Premix Ex Taq™ II (Tli RNaseH Plus) kit on QuantStudio 6 and 7 Flex real-time PCR systems (Thermo Fisher, USA). Reagents mentioned above were obtained from Takara (Japan). Primers for RT-qPCR were provided by Sangon Co. (Shanghai, China) and attached to the Appendix Table S1-2. The extracellular or plasma levels of Il-1β, Il-6, Tnf-α, BDNF, GDNF, and NGF were confirmed by ELISA using commercially available kits.

### Western Blotting

Clarified cell and tissue lysates were fractionated by SDS-PAGE using 8-15% gradient polyacrylamide gels and transferred onto a PVDF membrane. After blocking with TBST containing 5% non-fat milk (Bio-Rad, USA) for 1 h, the membranes were stained with primary antibodies in the diluted solution overnight at 4 °C. Primary antibodies used were: anti-MAP2 (1:1000), anti-NeuN (1:1000), anti-Synaptophysin (1:5000), anti-GFAP (1:5000), anti-C3/C3b/C3c (1:1000), anti-CHRNA7 (1:2500), anti-JAK2 (1:500), anti-STAT3 (1:2000), rabbit monoclonal anti-phosphorylated-JAK2 (1:5000, phospho Y1007+Y1008, Abcam, UK), rabbit monoclonal anti-phosphorylated-STAT3 (1:10000, phospho Y705, Abcam, UK), and mouse monoclonal anti-GAPDH (1:10000). Except for the indicated antibodies, other antibodies mentioned above were rabbit polyclonal antibodies purchased from Proteintech (USA). Following incubation with appropriate HRP-conjugated secondary antibodies (1:5000, Bioss, USA) for 1 h at room temperature, the membrane was treated with Clarity Western ECL blotting substrate (Bio-Rad, 1705060, USA). Bands were quantified using Fuji ImageJ software.

### Immunolabeling assay for primary astrocytes

Cells were fixed with 100% methanol for 20 min at −20°C and permeabilized with 0.25% Triton X-100 prepared in PBS for 10 min, followed by blockage with mixture buffer (1% w/v BSA, Sigma-Aldrich, USA; 22.52 mg/ml glycine in PBST). The cells were incubated with mouse monoclonal Cy3-conjugated anti-GFAP (1:400, not requiring secondary antibody, Abcam, UK), anti-C3/C3b/C3c (1:500), Alexa Fluor 488-conjugated α-bungarotoxin (5 μg/ml, Thermo Fisher Invitrogen, USA), anti-JAK2 (1:200), anti-STAT3 (1:100) for overnight at 4°C, followed by incubation with Alexa Fluor 488-conjugated goat anti-rabbit secondary antibody (1:500), goat anti-rabbit IgG H&L (Cy5, 1:2500, Abcam, UK) for 2 h in the dark at room temperature. DAPI (4 μg/mL, Thermo Fisher, USA) was used as nuclei stain. Stained cells were examined using a fluorescent microscope (E800 Nikon, Japan) connected to a color digital camera.

### Animals and treatment

The HIV-1 gp120 transgenic mice (gp120tg mice, 3 months old, 6 months old, 12 months old, 24 months old; *n*=10-15 per group) on SJL/BL6/129 background (cross between C57BL/6J female x Sv129 male) expressed gp120 in astrocytes under the control of a modified murine *Gfap* gene. The α7nAChR knockout mice (α7^-/-^ mice, B6.129S7-Chrna7tm1Bay/J) were purchased from the Jackson Laboratory. The gp120tg mice intercrossed with α7^-/-^ mice to generate α7^-/-^gp120tg mice. C57BL/6J black mice (12 months old) were used as wild-type controls. Genotypes of all mice were confirmed by PCR analysis of tail DNA with the following primer sequences: gp120 forward, 5’-GCGGGAGAATGATAATGGAG-3’; gp120 reverse, 5’-TATGGGAATTGGCTCAAAGG-3’; α7nAChR forward, 5’-TTCCTGGTCCTGCTGTGTTA-3’; α7nAChR wild-type reverse, 5’-ATCAGATGTTGCTGGCATGA-3’; α7nAChR knockout reverse, 5’-CCCTTTATAGATTCGCCCTTG-3’. All mice were bred in groups of 5 per cage and maintained in a pathogen-free animal facility with free access to autoclaved distilled water and standard chow under a constant temperature of 22 ± 1 °C on a 12 h light/dark cycle. For tryptophan supplement, mice were fed the control diet supplemented with 0.1% tryptophan (Sigma-Aldrich, USA) dissolved in sterile saline (0.9 % NaCl) for 1 month. All procedures involving animals were approved by the Experimental Animal Ethical Committee of Southern Medical University (ethics project identification code: 2015029).

### Morris water maze test and open field test

For the MWM test, mice used 4 unique geometric figures providing landmarks in the testing room to locate a submerged platform (1 cm below the white-opaque water surface) in a circular pool (1.2 m in diameter and 76 cm in height). Mice were allowed 60 s to locate the hidden platform in one maze quadrant under a randomized starting position for 4 trails per day over 5 consecutive days. Upon completion of training, mice were allowed to locate the removed platform for 60 s in the retrieval test. Performances were recorded with a camera suspended 250 cm above the center and analyzed by the image tracking system. Mice were evaluated in the open field test of anxiety and exploratory behavior. Briefly, mice were allowed to move unfetteredly in an enclosure and brightly lit square arena (40×40 cm) for 15 minutes on 2 consecutive days. Distance moved, and time spent in the central area were videotaped with a near-infrared camera positioned above the center of the arena. The central zone was 32 cm in diameter with 4 cm from the peripheral walls. An effective center entry was not deemed to have occurred until 90% or over of the mouse’s body had entered the designated center.

### Histopathology assessment

Upon anesthesia of mice by sodium pentobarbital (60 mg/kg, Sigma-Aldrich, USA), brains were dissected and fixed by immersion in 10% formalin solution overnight at room temperature. For viewing cellular and tissue structure detail, deparaffinized and rehydrated slides were counterstained with Mayers Hematoxylin and alcoholic-Eosin (H&E). In addition, Nissl-stained sections were labeled with 0.1% cresyl violet solution at 37 °C for 10 min after deparaffinization and hydration. Next, the slides were differentiated in 95% ethyl alcohol for 5 to 10 min, followed by incubation with 100% alcohol and xylene.

### Immunofluorescence and immunohistochemistry

For immunofluorescence, deparaffinized and rehydrated brain sections were stained with anti-MAP2 (1:500), anti-NeuN (1:500), Cy3-conjugated anti-GFAP (1:400), and anti-C3/C3b/C3c (1:500). Following PBS washes, slides were reacted with Alexa Fluor 488-labeled goat anti-rabbit antibody (1:1000) at room temperature for 1 h. The slides were washed and subsequently counterstained with DAPI (1.5 μg/mL, Thermo Fisher, USA). Five sections from the hippocampus were randomly selected, and the numbers of C3^+^/GFAP^+^ cells were counted. For immunohistochemistry, slides were reacted with anti-phosphorylated-JAK2 (1:100) and anti-phosphorylated-STAT3 (1:100) antibodies overnight at 4 °C in a humidity chamber. After rinsing with PBS, sections were stained with a biotinylated antibody for 60 min and then incubated with 3,3’-diaminobenzidine tetrahydrochloride (DAB) solution for 5 min. Subsequently, sections were counterstained with hematoxylin.

### Statistical analysis

Graphs and statistical significance are generated by GraphPad Prism 8 (version 8.3.0). The normality and homogeneity of variance were assessed by the *Shapiro-Wilk* test and *Levene’s* test, respectively. Data were analyzed using an unpaired Student’s *t* test (two-tailed) for two-group comparison. One-way ANOVA followed by *post hoc* Dunnett’s multiple comparisons was used to compare more than three groups when compared with the one indicated group.

## Acknowledgments

This project was financially supported by the National Natural Science Foundation of China (No. 82172259) and the Major Program of the Natural Science Foundation of Guangdong, China (No. 2017B030311017).

## The Paper Explained

### Problem

HIV-1 gp120 has long been considered a critical player in neuropathologies of HIV-associated neurocognitive disorder (HAND). It is putative that the generation of neurotoxic reactive astrocytes (A1 astrocytes) bears responsibility for neurodegeneration. Nevertheless, mechanisms facilitating A1 astrocyte activation and exhaustion of A2 astrocyte homeostatic capacities under HAND are not fully understood.

### Results

Our research reveals that the activation of A1 astrocytes is already evident in 6-month-old gp120tg mice, contributing to neuronal aberration and cognitive declines. Kynurenic acid (KYNA), an α7nAChR antagonist, promotes A1 to A2 astrocyte transformation via blockage of α7nAChR/JAK2/STAT3 cascades. Notably, signs of A1 to A2 astrocyte conversion are observed not only in α7-/-gp120tg mice but also in tryptophan-fed (the precursor of KYNA) gp120tg mice, and this phenotypic transformation rescues neuronal loss and behavioral deficits in aged gp120tg mice.

### Impact

Overall, our paper highlights the neuroprotective properties of KYNA in HAND or other neurologic insults involved in A1 astrocyte activation.

## Data availability

This study includes no data deposited in external repositories. The data that support the plots within this study are available from the corresponding author upon reasonable request.

## Expanded View Figure Legends

**Figure EV1 - Age-dependent neurological and cognitive declines in gp120tg mice.**

A H&E (scale bar: 100 μm) or Nissl body (scale bar: 10 μm) showed the histology of the hippocampus. Black arrows indicated a group of Nissl bodies (*n*=5 biologically independent animals).

B Quantification of MAP-2, NeuN, and Syn protein levels in the hippocampus of WT and gp120tg mice (*n*=5 mice per group).

C-F Latency (C) and total distance (D) to locate the submerged platform during 5 training days in the MWM test among 3-, 6-, 12-, and 24-month-old gp120tg mice. The behavior (E) and time spent (F) in the 60 s probe trial (*n*=10 mice per group).

G-J WT and gp120tg mice were allowed to explore undisturbed in the square open-field box for 15 min. Representative performance of mice during the open field test. Red dots reflected the starting position, and green dots depicted the ending position. Bars show the average time spent in the (H) central area, (I) distance traveled in the central zone, and (J) the number of times entering the center (*n*=8 mice per group).

Data information: In (B-F), data are presented mean ± SD. *P* values were analyzed by one-way ANOVA. In (H-I), *P* values were analyzed by Student’s *t* test. \**P* < 0.05, \*\**P* < 0.01, \*\*\**P* < 0.001, NS *P* > 0.05 (not significant) compared to WT mice. ^$^*P* < 0.05, ^$$^*P* < 0.01, ^$$$^*P* < 0.001 compared to 6-month-old gp120tg mice.

**Figure EV2** - **KYNA attenuates the activation of the α7nAChR/JAK2/STAT3 signaling pathway in gp120 induced astrocytes.**

A Quantification of GFAP and C3 protein levels by Western blot.

B Astrocytes were treated with gp120 at a wide range of concentrations from 1.5 pM to 1500 pM for 24 h. The expression of α7nAChR on the cell membrane is indicated by α-bungarotoxin binding images (scale bar: 100 μm).

C The α7nAChR protein levels in lysate from astrocytes exposed to increasing doses of gp120.

D Astrocytes were exposed to 150 pM gp120 for 12 h, then co-incubated with 5 or 25 μM KYNA for another 12 h. Immunofluorescence showing the effects of KYNA on α7nAChR expression (scale bar: 100 μm).

E JAK2-positive astrocytes (green).

F STAT3-positive cells (orange).

Data information: In (A-F), representative data are paresented as mean ± SD with *n* = 5 cultures per group; one-way ANOVA. *\*P* < 0.05, \*\**P* < 0.01, \*\*\**P* < 0.001, NS *P* > 0.05 (not significant) compared to 150 pM HIV-1 gp120 treated group.

**Figure EV3** - **KYNA administration reversed the effects of nicotine or PNU-282987 pretreatment in the phenotypic transformation of gp120-induced astrocytes.**

A-C Following 12 h treatment with 150 pM gp120, primary rat astrocytes were pretreated with 10 μM nicotine or 0.5 μM PNU-282987 for 1 h, then co-incubated with 25 μM KYNA for 11 h. The mRNA levels of A1-specific genes (A), A2-specific genes (B), and cytokines (C) in nicotine-treated astrocytes.

D-F Following 12 h treatment with 150 pM gp120, primary rat astrocytes were pretreated with 0.5 μM PNU-282987 for 1 h, then co-incubated with 25 μM KYNA for 11 h. The mRNA levels of A1-specific genes (D), A2-specific genes (E), and cytokines (F) in PNU-282987-stimulated astrocytes.

Data information: Values are shown in the box and whisker plot where the line in the box corresponds to the median, one-way ANOVA, *n* = 5 replicates per group. \**P* < 0.05, \*\**P* < 0.01, \*\*\**P* < 0.001, NS *P* > 0.05 (not significant) compared to 150 pM gp120 treated group. ^$^*P* < 0.05, ^$$^*P* < 0.01, ^$$$^*P* < 0.001 compared to gp120+nicotine group or gp120+PNU-282987 group.

